# NIX-associated mitochondrial remodeling contributes to esophageal epithelial differentiation

**DOI:** 10.64898/2026.09.17.752311

**Authors:** No’ad Shanas, Aissata Balde, Annie D. Fuller, Jazmyne L. Jackson, Abigail J. Staub, John M. Crespo, William Nazario-Lugo, Alexander Tufano, Nicholas M. McDonald, Andres J. Klein-Szanto, Kathy Q. Cai, Melanie Ruffner, Amanda B. Muir, Zachary Wilmer Reichenbach, Kelly A. Whelan

## Abstract

Mitochondria are increasingly recognized as regulators of cellular differentiation, but their role in esophageal epithelial homeostasis remains poorly understood. Here, we investigated mitochondrial remodeling during esophageal epithelial differentiation and whether mitochondrial depletion contributes to the acquisition of the differentiated phenotype. Mitochondrial abundance was assessed across normal human esophageal epithelium and in non-transformed immortalized human esophageal epithelial cells (EPC2-hTERT) using two differentiation models. Mitochondrial architecture was quantified, and the functional role of mitochondrial abundance was examined using doxycycline-inducible depletion of transcription factor A, mitochondrial (TFAM). Mitochondrial abundance progressively decreased from basal to superficial compartments of normal human esophageal epithelium and during *in vitro* differentiation, accompanied by fragmentation and remodeling of the mitochondrial network. TFAM depletion reduced mitochondrial abundance and increased squamous differentiation markers, indicating that mitochondrial depletion was sufficient to promote differentiation. Analysis of candidate mitochondrial clearance pathways identified Bcl2-interacting protein 3-like (*BNIP3L*)/NIX (NIP3-like protein X) as preferentially associated with differentiated epithelial cells and induced during differentiation. NIX was associated with mitochondria and increased as mitochondrial abundance declined. NIX depletion prevented differentiation-associated mitochondrial depletion and attenuated differentiation marker induction. In human esophageal epithelium, NIX expression increased across the basal-to-suprabasal compartment before declining superficially. *BNIP3L* expression was reduced in active eosinophilic esophagitis (EoE) and increased following corticosteroid-associated remission. Interleukin-13 similarly reduced *BNIP3L* expression and suppressed epithelial differentiation in EPC2-hTERT cells. These findings identify NIX-associated mitochondrial remodeling as an important component of esophageal epithelial differentiation and suggest that disruption of mitochondrial quality control may contribute to impaired epithelial differentiation in EoE.

**New and Noteworthy:** Although impaired esophageal epithelial differentiation is associated with various esophageal diseases, mechanisms regulating differentiation under homeostasis remain incompletely understood. We show that mitochondria are progressively cleared during esophageal epithelial differentiation and that mitochondrial depletion is sufficient to promote differentiation. BNIP3L/NIX regulates mitochondrial clearance during this process. Moreover, BNIP3L expression is reduced in active eosinophilic esophagitis (EoE) and restored with remission, linking disrupted mitochondrial quality control to epithelial dysfunction in EoE.

## Introduction

Esophageal squamous epithelium is a first-line barrier against luminal insults, including allergens, pathogens, and carcinogens. To maintain this barrier, the epithelium undergoes continuous renewal through a spatially organized differentiation program in which proliferative basal cells give rise to suprabasal cells that progressively exit the cell cycle and acquire a differentiated phenotype toward the luminal surface. This process is characterized by coordinated changes in epithelial structure and gene expression, including induction of differentiation-associated proteins such as involucrin (IVL) and cytokeratin 13 (CK13) (1). Disruption of this differentiation program is a feature of common esophageal diseases, including eosinophilic esophagitis (EoE) and esophageal cancer (2, 3). Defining the cellular mechanisms that establish and maintain esophageal epithelial differentiation may therefore provide insight into both normal tissue homeostasis and disease pathogenesis.

Mitochondria are increasingly recognized as dynamic regulators of cell state and differentiation (4–6). In several epithelial and non-epithelial lineages, differentiation is accompanied by changes in mitochondrial metabolism, abundance, and architecture. In some contexts, selective mitochondrial clearance through mitophagy is required for cells to acquire the differentiated phenotype (7, 8). Conversely, mitochondrial biogenesis and the expansion of mitochondrial mass support differentiation in other tissues, including the intestinal epithelium (9). Thus, mitochondrial remodeling can be coupled to differentiation, but the nature and functional consequences of this remodeling appear to be highly dependent on cellular context.

Despite the importance of mitochondrial regulation in other differentiating tissues, little is known about mitochondrial remodeling during esophageal epithelial differentiation. Recent studies have identified alterations in mitochondrial abundance and function in EoE (10–12), raising the possibility that mitochondrial dysregulation contributes to epithelial remodeling in this disease. However, whether mitochondrial abundance and architecture are spatially regulated across the normal esophageal epithelium, whether mitochondrial remodeling contributes to the acquisition of the differentiated state, and how mitochondria are cleared during esophageal epithelial maturation remain unknown.

Here, we investigated mitochondrial remodeling during esophageal epithelial differentiation and determined whether mitochondrial clearance contributes to the acquisition of the differentiated phenotype. We demonstrate that mitochondrial abundance progressively decreases as esophageal epithelial cells differentiate and that this reduction is accompanied by remodeling and fragmentation of the mitochondrial network. Experimentally induced mitochondrial depletion was sufficient to promote features of epithelial differentiation, suggesting that mitochondrial reduction is not simply a consequence of differentiation. We further identified Bcl2-interacting protein 3-like (*BNIP3L*)/NIX as a candidate mediator of differentiation-associated mitochondrial clearance. NIX was induced during esophageal epithelial differentiation, localized to mitochondria, and was required for efficient mitochondrial clearance and the induction of differentiation-associated markers. Finally, we found that *BNIP3L* expression was reduced in active EoE and in esophageal epithelial cells treated with the EoE-relevant cytokine interleukin-13 (IL-13) (13,14), supporting a potential link between NIX-associated mitochondrial remodeling and the altered epithelial state observed in human disease.

## Materials and methods

### Human Subjects

All biological specimens were obtained from patients in collaboration with Temple University Hospital (TUH) under Dr. Reichenbach’s IRB-approved protocol (#27291). Human subjects underwent endoscopic biopsy collection via esophagogastroduodenoscopy (EGD) during routine clinical care at TUH. At the time of diagnostic EGD, pinch biopsies were obtained for clinical evaluation. Additional biopsy specimens were collected for research. Written informed consent was obtained from each subject. All subjects reported symptoms warranting EGD but demonstrated no endoscopic or histological abnormalities in the esophagus. Subjects with a history of inflammatory bowel disease, celiac disease, esophageal surgery, malignancy, or treatment with radiation were excluded from recruitment. All subjects were confirmed to have normal esophageal pathology following review by the study pathologist (AJK-S). Demographic and clinical information on human subjects whose esophageal biopsies were evaluated in the current study is provided in supplementary **Table 1**.

**Table 1:**
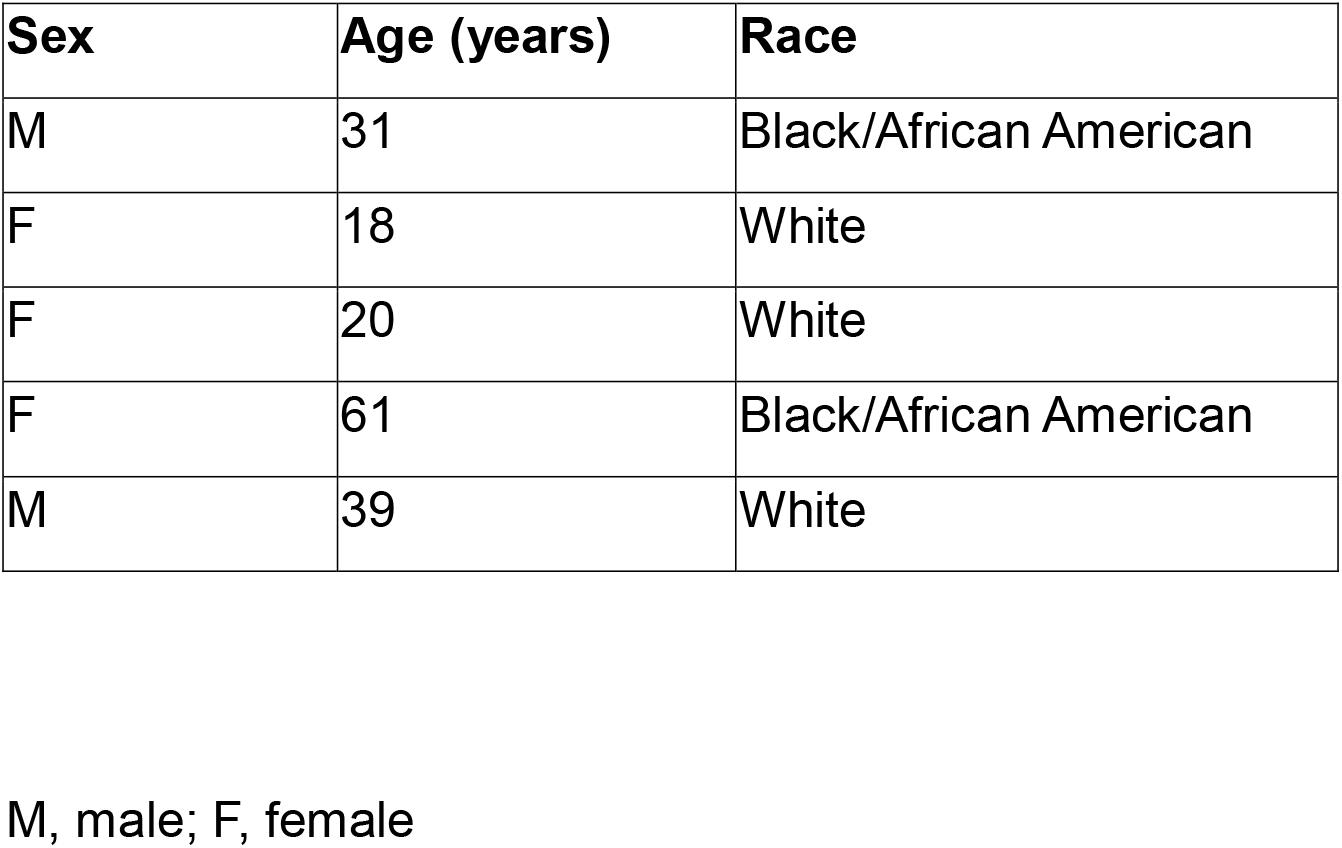
Human subject demographic information.

### Slide Preparation and Immunohistochemistry (IHC)

Whole esophagi were dissected, fixed in 10% neutral buffered formalin for 12 hours at room temperature, then dehydrated and embedded in paraffin. Hematoxylin and eosin (H&E) stained sections were used for morphological evaluation, and 5 µm-thick unstained sections were used for IHC studies. Immunohistochemical staining was performed on a Ventana Discovery Ultra automated staining instrument (Ventana Medical Systems) using Ventana reagents according to the manufacturer’s instructions. Briefly, slides were deparaffinized using EZ Prep solution (950–102) for 16 min at 72 °C. Epitope retrieval was accomplished with CC1 solution (EDTA, pH 9.0; 950–224) at high temperature (95–100 °C) for 32 min. Mouse primary antibodies against translocase of outer mitochondrial membrane 20 (TOM20) and oxidative phosphorylation (OXPHOS) cocktail (**Table 2**) were titrated in TBS antibody diluent and loaded into user-fillable dispensers for use on the automated stainer. Immune complexes were detected using Ventana OmniMap anti-mouse detection kit (760-4310) and developed using Ventana ChromMap DAB detection kit (760-159) according to the manufacturer’s instructions. Slides were counterstained with hematoxylin II for 8 min followed by Bluing reagent for 4 min. Slides were then dehydrated with an ethanol series, cleared in xylene, and mounted. All stained sections were digitized as whole-slide images using a NanoZoomer S60 digital slide scanner (Hamamatsu Photonics, Hamamatsu, Japan) at 40X magnification (0.23 µm/pixel) under brightfield acquisition settings. Images were reviewed for focus quality, tissue integrity, and staining uniformity before downstream analysis. Slides for linear integrated density quantification were imaged using a Leica DM 1000 LED microscope at a 20X objective lens. To quantify protein expression in human esophageal biopsies, IHC images were analyzed using ImageJ software. Images underwent color deconvolution to isolate the DAB signal. To assess TOM20 and OXPHOS protein expression in human esophageal biopsies, 5 regions of interest (ROIs) were selected from the basal compartment (20% of the epithelium located closest to the underlying stroma) and 5 ROIs from the superficial compartment (20% of the epithelium located closest to the lumen). To assess expression gradients across the epithelium, five straight lines were traced per image, extending across the basal-to-superficial epithelial axis, to measure integrated densities. To account for variations in epithelial thickness across sections and cellular compartments, each spatial profile was normalized by binning the data into 75 equal points. The density values from the five lines were then averaged, normalized to the maximum intensity (expressed as a percentage), and smoothed using a second-order polynomial with a 4-neighbor window. This processing enabled the clear visualization of relative protein expression changes from the basal to the superficial layers.

**Table 2:** List of antibodies.

| Antibody | Source | Catalog number | Application<br>(dilution) |
| --- | --- | --- | --- |
| OXPHOS | ABCAM | 110411 | IHC |
| TOM20 | Santa Cruz | 17764 | IHC |
| TFAM | Sigma | 7495S | IB: 1:1000 |
| Parkin | Cell Signaling | 4211S | IB: 1:500 |
| CK13 | Fisher | NC1907584 | IB: 1:1000 |
| TOM20 | ABCAM | ab289670 | IB: 1:1000<br>ICC: 1:100 |
| Actin | Invitrogen | MA1-744 | IB: 1:5000 |
| NIX | Cell Signaling | 12396S | IB: 1:1000<br>ICC: 1:100 |
| Alexa Fluor 488 | Biolegend | 405319 | ICC: 1:1000 |
| Alexa Fluor 555 | Fisher | A21428 | ICC: 1:1000 |
| Alexa Fluor 647 | Fisher | A21247 | ICC: 1:1000 |
| anti-Rabbit HRP | Fisher | PI31460 | IB: 1:10,000 |
| anti-Mouse HRP | Fisher | 50195914 | IB: 1:10,000 |
| anti-Rat HRP | Invitrogen | PI31470 | IB: 1:10,000 |

IB, immunoblot; IHC; immunohistochemistry; ICC, immunocytochemistry.

### Cell Culture

The non-transformed immortalized normal esophageal keratinocyte cell line EPC2-hTERT (15) was provided as a generous gift from Drs. Anil Rustgi and Hiroshi Nakagawa (Columbia University). Cells were cultured in keratinocyte serum-free medium (KSFM; Gibco, 17005042) supplemented with recombinant epidermal growth factor (1 ng/mL), bovine pituitary extract (50 mg/mL), and penicillin/streptomycin (1% v/v, Gibco, 15140-122), as previously described (10). For calcium (Ca^2+^)-induced differentiation, EPC2-hTERT cells were plated in 6-well plates (80,000 cells per well) and incubated for 48 hours in KSFM (0.09 mM Ca^2+^), then the medium was changed to fresh KSFM with 1.8 mM Ca^2+^ (16, 17). Media was changed every 48 hours for a total of 3-7 days. For confluence-induced differentiation, EPC2-hTERT cells were plated at 150,000 cells per well in 6-well plates and maintained in the same well for 3 weeks, with media refreshed every 2-3 days. Cells plated in 6-well plates at 60,000 cells per well were used as low-confluence controls. For interleukin (IL)-13 stimulation, EPC2-hTERT cells were plated in 6-well plates (30,000 cells per well) 24 hours before IL-13 (Bio-Techne, 213-ILB) at a final concentration of 10 ng/mL. Media was changed every 48 hours for a total of 7 days.

### Genetic Depletion

To deplete TFAM in esophageal keratinocytes, EPC2-hTERT cells with doxycycline (DOX)-inducible expression of short hairpin RNA (shRNA) targeting TFAM (Horizon Discovery, V3SH7669224733763) were generated by lentiviral transduction as previously described (10). EPC2-hTERT cells expressing non-targeting (NT) shRNA (Horizon Discovery, VSC11653) served as controls. 60,000 cells were plated in a 6-well plate and were treated with DOX (0.5 µg/mL) for 7 days to initiate TFAM depletion. To achieve NIX knockdown in esophageal keratinocytes, small interfering RNA (siRNA) oligonucleotides directed against NIX (Fisher, 4392420) or a NT control pool (Fisher, 4390843) were prepared in 500 μL of Opti-MEM reduced serum medium (Gibco, 31985088) to yield a final concentration of 10 nM per well in a 6-well plate. Next, 5 μL of Lipofectamine™ RNAiMAX transfection reagent (Thermo Fisher Scientific, 13778150) was added to each well. Plates were incubated for 20 minutes at room temperature before introducing 300,000 EPC2-hTERT cells suspended in 2 mL of antibiotic-free KSFM to each well. Cells were then incubated for 72 hours post-transfection before use in downstream experiments.

### Quantitative Reverse Transcription Polymerase Chain Reaction (qRT-PCR)

Total RNA was extracted using the RNeasy Mini Kit (Qiagen, 74106) according to the manufacturer’s instructions, prior to qRT-PCR analysis. RNA concentration was measured using the Qubit™ RNA HS Assay Kit (Invitrogen, Q32852). Reverse transcription of extracted RNA was performed utilizing the High-Capacity cDNA Reverse Transcription Kit (Thermo Fisher Scientific, 4368814), with subsequent target gene quantification achieved via real-time qRT-PCR using the PowerUp™ SYBR™ Green Master Mix (Thermo Fisher Scientific, A25743). Primers for β-Actin (*ACTB*), BCL2/adenovirus E1B 19 kDa interacting protein 3 *(BNIP3), BNIP3L, KRT13,* cytokeratin 14 (*KRT14*), *IVL*, *TFAM*, and Parkin (*PARK2*) were used. The relative expression of each gene, normalized to β-Actin, was calculated using the ΔΔCt method. All primer sequences are listed in **Table 3**.

**Table 3:**
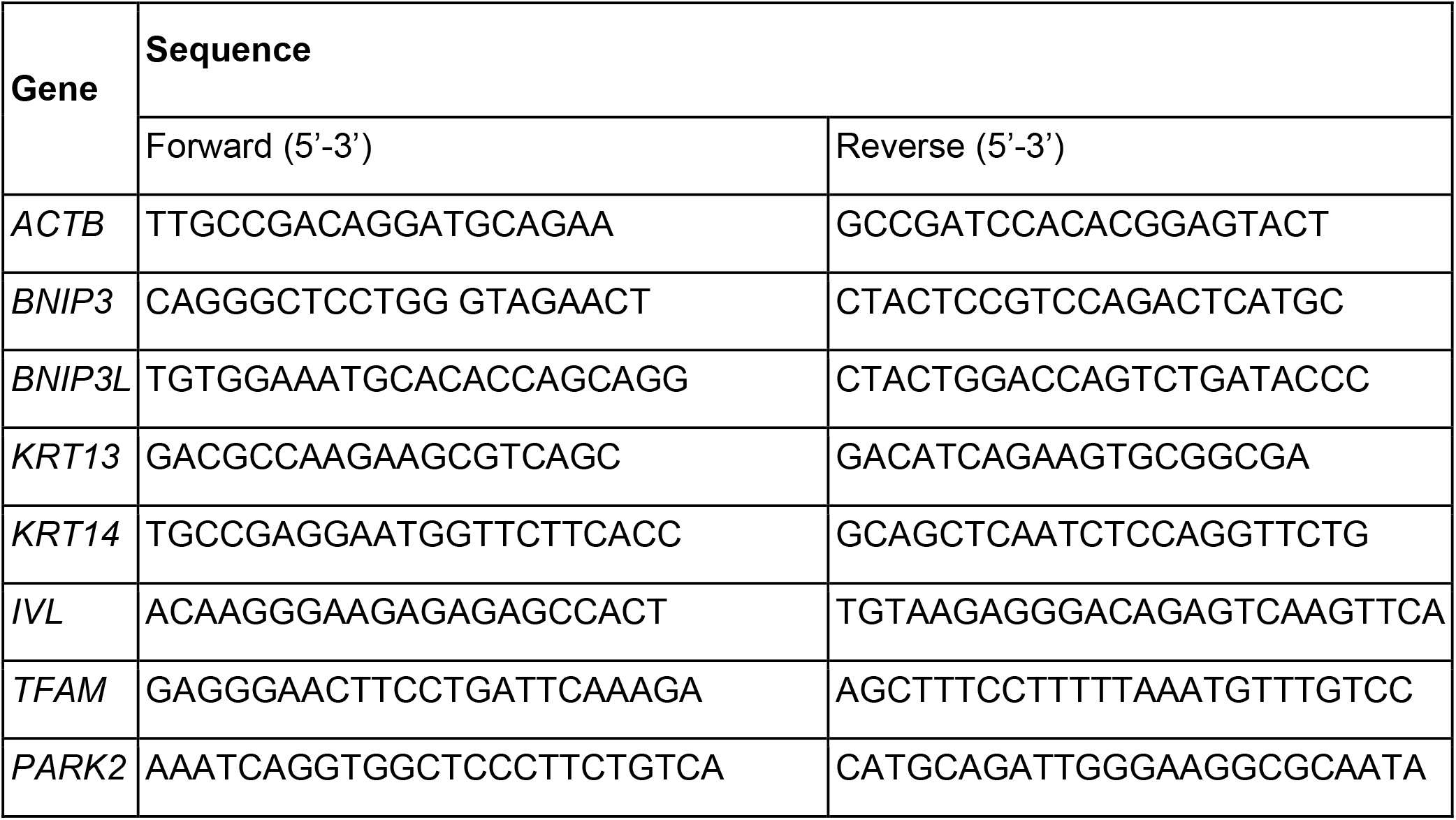
List of primers qRT-PCR primers.

### Immunocytochemistry

Esophageal keratinocytes were plated in 6-well plates with 25 mm collagen-coated coverslips overnight. Cells were fixed with 4% paraformaldehyde, washed once with 1X PBS, then permeabilized with 1X PBST (0.15% Triton X in 1X PBS) for 10 minutes. Cells were then blocked with 5% bovine serum albumin and 300 mM glycine in 1X PBST for 1 hour, and incubated overnight at 4°C with the primary antibodies listed in **Table 2**. Following 1X PBST washes, cells were incubated with secondary antibodies (**Table 2**) for 1 hour at room temperature, protected from light. Nuclei were then counterstained with 2 µM DAPI (Fisher, 62248), mounted using ProLong Gold Antifade (Fisher, P10144), and stored at 4°C prior to imaging.

### High Resolution Confocal Microscopy

Immunocytochemistry-fixed and stained slides were visualized using a Leica SP8 Laser Scanning Microscope equipped with two photomultiplier tubes and one hybrid detector, in addition to 405, 488, 522, and 638 nm lasers. Slides were imaged using a 63X oil objective (1.4 NA) and type F Leica microscope immersion oil (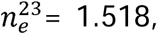 Thomas Scientific, 22A00L992), using LAS X Life Science Microscope Software. Scientific Volume Imaging Nyquist Calculator (https://svi.nl/Nyquist-Calculator) was used to determine the appropriate image voxel parameters. Images were captured at 72.22 nm × 72.22 nm with a 160 nm z-step size, 3-frame averaging, and a pinhole of 0.75 Airy Unit. Prior to downstream quantification and analysis, raw images were deconvolved using Huygens Essentials Software, using Classic Maximum Likelihood Estimation, 45 iterations (classical iterations mode), p<0.001, and slice-by-slice module option.

### Mitochondrial Network Analysis

Images were quantified using ImageJ software with the Mitochondria Analyzer plug-in previously described (18), with adaptive thresholding optimized at radius=2, C=4. The complete source code is publicly deposited on GitHub (https://github.com/AhsenChaudhry/Mitochondria-Analyzer).

### Live-Cell Imaging

To quantify mitochondrial mass, cells were incubated with 200 nM MitoTracker Green (Invitrogen, M7514) in KSFM for 30 min at 37°C in a 5% CO2 atmosphere. Following three washes with 1X PBS, live-cell imaging was conducted under standard culture conditions (37°C, 5% CO2), in KSFM, using an IncuCyte S3 Live-Cell Analysis System equipped with an S3/SX1 Optical Module (Sartorius). Phase-contrast and green fluorescence images were acquired at 20X magnification with a 400-ms exposure time per channel.

### Quantification of Live and Fixed Cell Fluorescence

Calibrated Incucyte or deconvolved confocal images were uploaded to ImageJ for quantification. For MitoTracker green assay, 10 cells were selected from each image field (3 images per trial, total of 90 cells per treatment group). Furthermore, for high-resolution confocal imaging quantification, individual cell images (25 images per trial, total of 75 cells per treatment group) were utilized for the analysis. To quantify fluorescence intensity, corrected total cell fluorescence (CTCF) was used (19, 20) and calculated with the following formula:

CTCF= Integrated density of ROI – (area of ROI x mean integrated density of background)

### Immunoblotting

Whole cell lysates from esophageal keratinocyte cultures were subjected to immunoblotting as previously described (10, 21). A list of antibodies used and their dilutions is provided in **Table 2**. Targeted proteins were visualized with chemiluminescence detection reagents (ProSignal Femto ECL Reagent, 20-302) and imaged on an iBright imaging system (Invitrogen).

### Publicly Available Data

To assess the expression of mitochondrial markers and mitophagy mediators in esophageal biopsies, the Human Protein Atlas (proteinatlas.org) was accessed (22). Specific gene expression profiles and representative immunohistochemical images for TOM20, translocase of inner mitochondrial membrane 23 (TIM23), mitochondrial NADH dehydrogenase subunit 3 (MTND3), mitochondrially encoded cytochrome C oxidase 1 (MTCO1), and NIX were evaluated. Image and data credit: Human Protein Atlas; specific images and data are available at proteinatlas.org (https://www.proteinatlas.org/ENSG00000104765-BNIP3L/tissue/esophagus; https://www.proteinatlas.org/ENSG00000173726-TOMM20/tissue/esophagus; https://www.proteinatlas.org/ENSG00000265354-TIMM23/tissue/esophagus; https://www.proteinatlas.org/ENSG00000198840-MTND3/tissue/esophagus; https://www.proteinatlas.org/ENSG00000198804-MT-CO1/tissue/esophagus)

To evaluate the expression of mitophagy mediators across distinct cellular compartments of the normal human esophagus, publicly available single-cell RNA-sequencing data (23) were downloaded from Tissue Stability Cell Atlas and analyzed using Seurat package in RStudio. Epithelial cell populations were subsetted and re-annotated to reflect sequential stratification stages (Basal, Suprabasal, Stratified, and Superficial), with cell states validated via canonical marker expression: DNA topoisomerase II alpha (*TOP2A*), *KRT14*, *IVL*, and cornulin (*CRNN*). Uniform Manifold Approximation and Projection (UMAP) feature maps, dot plots, and violin plots were generated to visualize expression profiles of mitophagy-related targets (*BNIP3L, BNIP3*, *IVL*, *PARK2*). Pairwise statistical comparisons between the basal compartment and suprabasal, stratified, or superficial layers were performed using a Wilcoxon rank-sum test.

Data from EPC2-hTERT cells grown in air-liquid interface (ALI) cultures or submerged monolayer cultures were acquired from the EGIDExpress database (https://egidexpress.research.cchmc.org/data/). In this study, EPC2-hTERT cells were grown continuously submerged for 8 days and served as the undifferentiated baseline control. These were compared with differentiated cells transitioned to the ALI for an additional 6 days to assess transcriptional changes using RNA sequencing (24).

Data for the comparison of gene expression in esophageal biopsies from histologically normal human subjects, EoE patients with active disease, and EoE patients treated with fluticasone were obtained from EGIDExpress. Briefly, distal esophageal biopsies were obtained from patients who had normal histological sections (control), chronic esophagitis (excluded from the current study), active EoE (≥24 eosinophils per high-power field), EoE patients who responded to fluticasone therapy, and EoE patients who did not respond to fluticasone treatment (the latter group was not included in the current study (25–27).

### Statistical Analysis

Descriptive statistics are presented as mean ± standard error of the mean (SEM). For Gaussian-distributed data, Student’s t-test or Welch’s t-test was used for two-group comparisons, and one-way analysis of variance (ANOVA) with Dunnett’s post hoc test was used for comparisons with more than 2 groups. For lognormally distributed data, unpaired lognormal ANOVA was used with Tukey’s post hoc test.

To assess the trial reproducibility of the lognormally distributed data, each trial was log-transformed using the following formula: “log(CTCF value)”. For sphericity measurement, the formula was adjusted to “log(CTCF value)+1” to eliminate negative data (for easier visualization). Average CTCF was calculated per trial, and trials were graphed to visualize changes. Statistics on the transformed data were done using one-way ANOVA with Tukey’s post hoc test. All statistics were performed using GraphPad Prism version 11.0.1 (90) for Windows (GraphPad Software, San Diego, California, USA). p<0.05 was considered statistically significant. For all data, the following indicators of significance are used: *p<0.05; **p<0.01; ***p<0.001; ****p<0.0001.

## Results

### Mitochondrial abundance is spatially patterned across the esophageal epithelial differentiation gradient

We first examined the distribution of mitochondria across the differentiation gradient of the normal human esophageal epithelium. Using IHC analysis of normal human esophageal tissue, we observed enriched expression of the mitochondrial structural proteins TOM20 and TIM23, as well as the mitochondrial-encoded oxidative phosphorylation (OXPHOS) proteins MTCO1 and MTND3 in basal epithelial cells (**Figure 1A**). Quantitative analysis confirmed significantly greater TOM20 and OXPHOS expression in the basal compartment compared to the superficial epithelial compartments (**Figure 1B-C; Supplementary Figure 1**). These findings reveal a distinct spatial gradient of mitochondrial abundance across the normal esophageal epithelium, with mitochondrial protein expression progressively decreasing as the epithelial cells differentiate toward the luminal surface.

**Figure 1.**
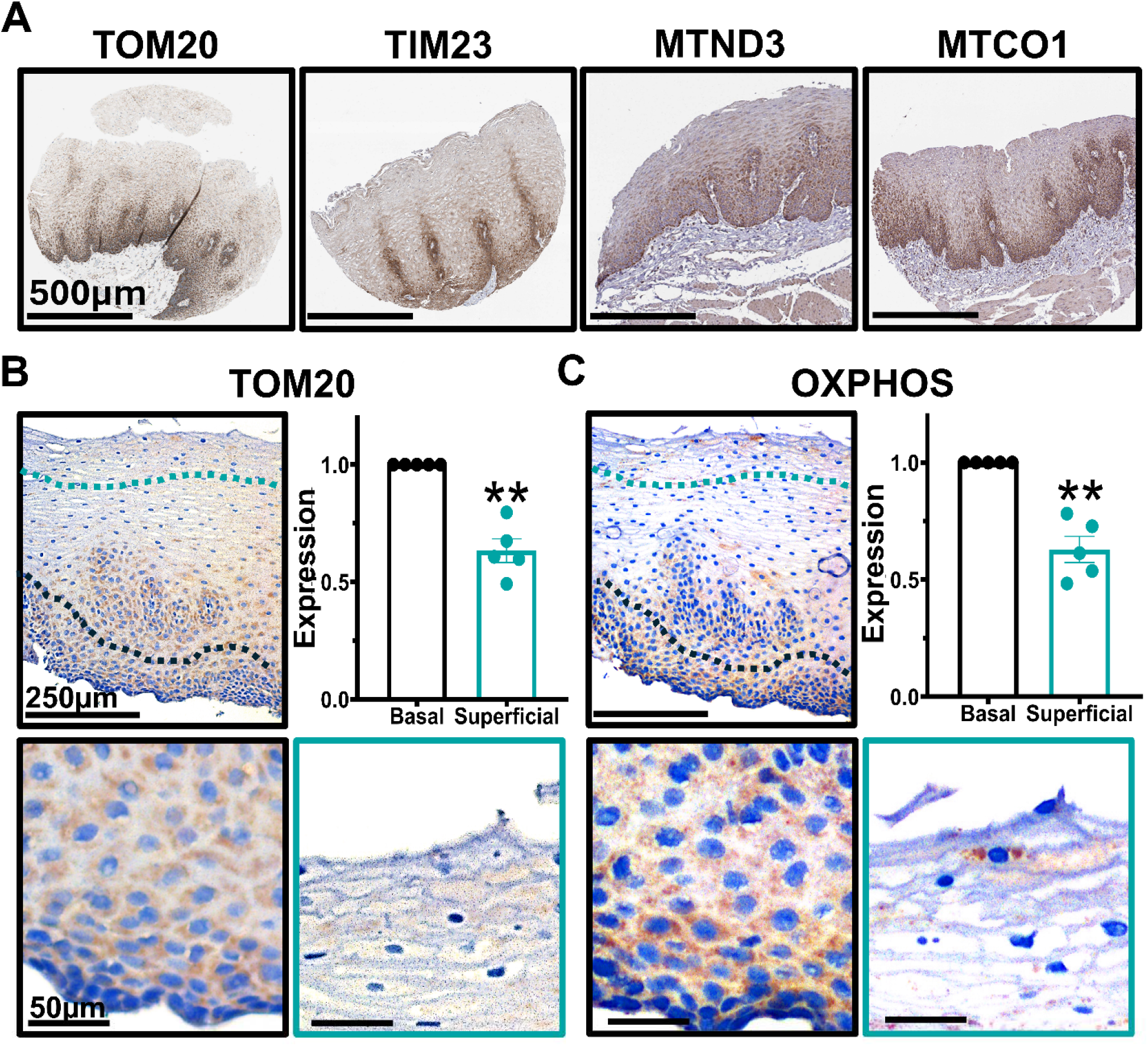
Mitochondrial abundance decreases along the basal-to-superficial axis of the esophageal epithelium. (**A**) Representative immunohistochemical (IHC) staining of human esophageal squamous epithelium from Human Protein Atlas for the mitochondrial membrane proteins TOMM20 and TIM23, and OXPHOS complex subunits MTCO1 and MTND3. (**B-C**) Representative IHC staining of human esophageal biopsies for (**B**) TOM20 and (**C**) OXPHOS. Staining intensity was quantified in the basal compartment (lower 20% of the epithelium; below black dashed line) and superficial compartment (upper 20%; above cyan dashed line). Scale bars, 500 μm (**A**), 250 μm and 50 μm (**B-C**). Each dot represents an individual human subject; n=5; **p<0.01 by Welch’s t-test. Data are presented as mean ± SEM.

### Esophageal epithelial differentiation is accompanied by mitochondrial depletion

We next asked whether the spatial gradient of mitochondrial abundance observed in human esophageal epithelium is recapitulated during epithelial differentiation *in vitro*. We investigated this by culturing EPC2-hTERT cells in standard low-calcium medium or transferring them to a high-calcium differentiation medium (1.8 mM Ca²⁺) for 3 or 7 days (**Figure 2A**). Calcium-induced differentiation was confirmed by the progressive induction of the differentiation markers *KRT13* and *IVL,* along with a reduction in the basal marker *KRT14* (**Figure 2B**). In parallel, MitoTracker Green staining demonstrated a significant reduction in fluorescence during differentiation (**Figure 2C-D**). To determine whether this relationship extended beyond calcium-induced differentiation, we used a second model in which increased cell density promotes epithelial differentiation. High confluence induced the mRNA expression of *IVL* and *KRT13,* and reduced *KRT14* expression (**Figure 2E-F)**. Furthermore, TOM20 protein abundance was diminished alongside the induction of CK13 protein (**Figure 2G-H**). Together, these findings demonstrate that mitochondrial abundance depletion accompanies esophageal epithelial differentiation across two independent *in vitro* models.

**Figure 2.**
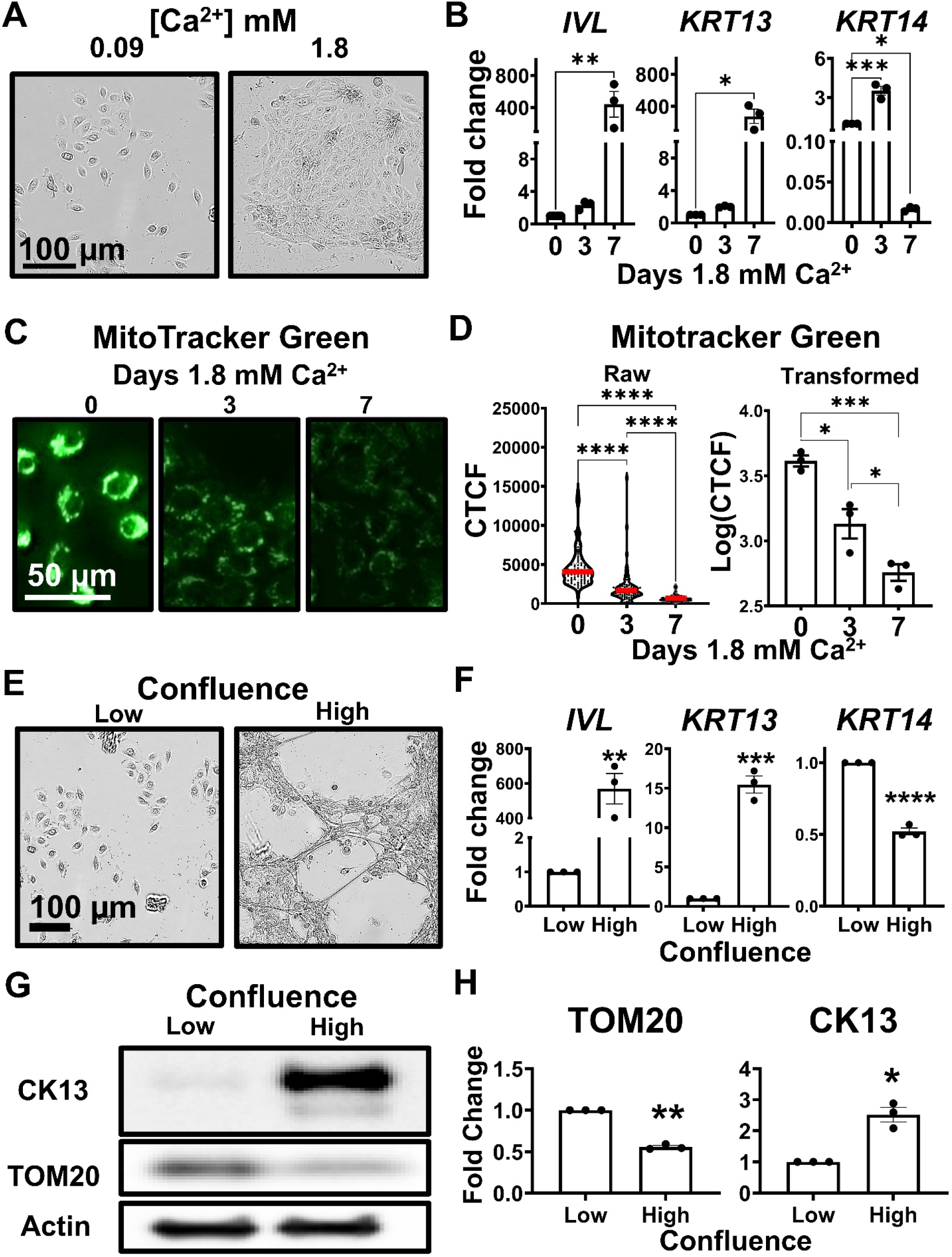
Esophageal epithelial differentiation is associated with reduced mitochondrial abundance *in vitro*. (**A-D**) EPC2-hTERT cells were cultured in low calcium maintenance medium (0.09 mM) and then transitioned to high calcium medium (1.8 mM) for 3 or 7 days. (**A**) Representative images of EPC2-hTERT cells cultured for 7 days. Scale bars, 100 μm. (**B**) Relative expression of differentiation markers *IVL* and *KRT13* and basal marker *KRT14* measured by qRT-PCR. Each dot represents an independent experiment; n=3; *p<0.05; **p<0.01; ***p<0.001 by one-way ANOVA with Tukey’s post hoc test. (**C**) Representative images of EPC2-hTERT cells stained with MitoTracker Green following exposure to 1.8 mM Ca²⁺ for 0, 3, or 7 days. Scale bars, 50 μm. (**D**) Quantification of MitoTracker Green fluorescence as corrected total cell fluorescence (CTCF). Raw single-cell measurements are shown on the left; each dot represents an individual cell; red line indicates mean. Data were obtained from three independent experiments. Trial-level transformed values are shown on the right; each dot represents an independent experiment; *p<0.05; ***p<0.001; ****p<0.0001 by lognormal Brown-Forsythe and Welch’s ANOVA tests (raw data) or one-way ANOVA with Tukey’s post hoc test (transformed data). (**E-H**) EPC2-hTERT cells were cultured at low or high confluence. (**E**) Representative images. Scale bars, 100 μm. (**F**) Relative expression of differentiation markers *IVL* and *KRT13* measured by qRT-PCR. Each dot represents an independent experiment; n=3; **p<0.01; ***p<0.001; ****p<0.0001 by Welch’s t-test. (**G**) Representative immunoblot showing CK13 and TOM20 expression. Actin was used as a loading control. (**H**) Densitometric quantification of CK13 and TOM20 protein expression. Each dot represents an independent experiment; n=3; *p<0.05; **p<0.01 by Welch’s t-test. Data are presented as mean ± SEM of 3 independent trials.

### Mitochondrial depletion is sufficient to promote esophageal epithelial differentiation

The association between mitochondrial depletion and epithelial differentiation raised the possibility that mitochondrial loss may contribute directly to the acquisition of the differentiated phenotype. To test this, we used a DOX-inducible shRNA approach to deplete TFAM, a key regulator of mitochondrial maintenance, in EPC2-hTERT cells. We confirmed the DOX-induced TFAM depletion at both the transcript and protein levels (**Figure 3A-C**). TFAM depletion significantly reduced TOM20 protein, consistent with reduced mitochondrial abundance. Notably, TFAM depletion was accompanied by increased expression of the differentiation markers *IVL* and *KRT13* at the transcript level (**Figure 3A)** and increased CK13 protein expression (**Figure 3B-C**). Thus, experimentally reducing mitochondrial abundance was sufficient to promote features of esophageal epithelial differentiation in the absence of an exogenous differentiation stimulus.

**Figure 3.**
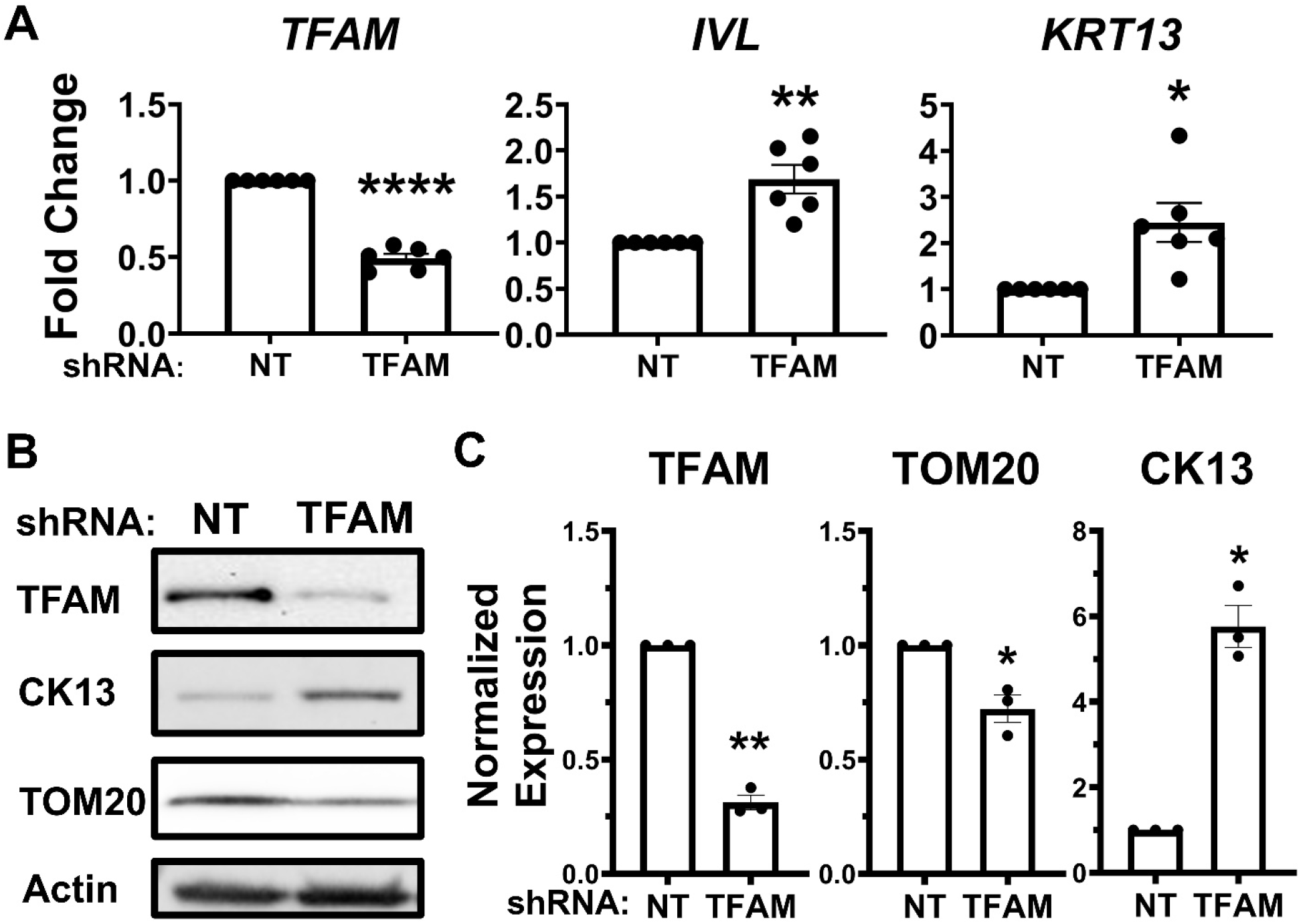
Mitochondrial depletion is sufficient to promote esophageal epithelial differentiation. EPC2-hTERT cells stably expressing doxycycline (DOX)-inducible shRNA targeting TFAM or a non-targeting control (NT) shRNA were treated with DOX for 7 days. (**A**) Relative expression of *TFAM*, *IVL*, and *KRT13* measured by qRT-PCR; n*=*6; *p<0.05; **p<0.01; ****p<0.0001 by Welch’s t-test. (**B**) Representative immunoblot showing TFAM, TOM20, and CK13 protein expression. Actin was used as a loading control. (**C**) Densitometric quantification of TFAM, TOM20, and CK13 protein expression. Each dot represents an independent experiment; n=3; *p<0.05; **p<0.01 by Welch’s t-test. Data are presented as mean ± SEM of independent trials.

### Esophageal epithelial differentiation is accompanied by mitochondrial network remodeling and fragmentation

Having established that epithelial differentiation is associated with mitochondrial clearance, we investigated whether it also alters mitochondrial network architecture. EPC2-hTERT cells were differentiated with 1.8 mM Ca²⁺ for 0, 3, or 7 days and stained for TOM20 to visualize the mitochondrial network. We then used high-resolution confocal microscopy, followed by automated three-dimensional image segmentation and morphometric analysis, to quantify mitochondrial architecture (**Figure 4A-B**). Differentiation was accompanied by a progressive loss of mitochondrial network connectivity, with mitochondria becoming increasingly discrete and fragmented by days 3 and 7 (**Figure 4C**). Although the number of individual mitochondria did not significantly change during differentiation, total mitochondrial volume and mean mitochondrial volume decreased significantly (**Figure 4D-F**). In parallel, mitochondrial sphericity increased and the number of branches per mitochondrion decreased (**Figure 4G-H**). Together, these findings demonstrate that esophageal epithelial differentiation is accompanied by substantial remodeling of mitochondrial architecture, characterized by reduced mitochondrial volume, increased sphericity, and loss of network branching.

**Figure 4.**
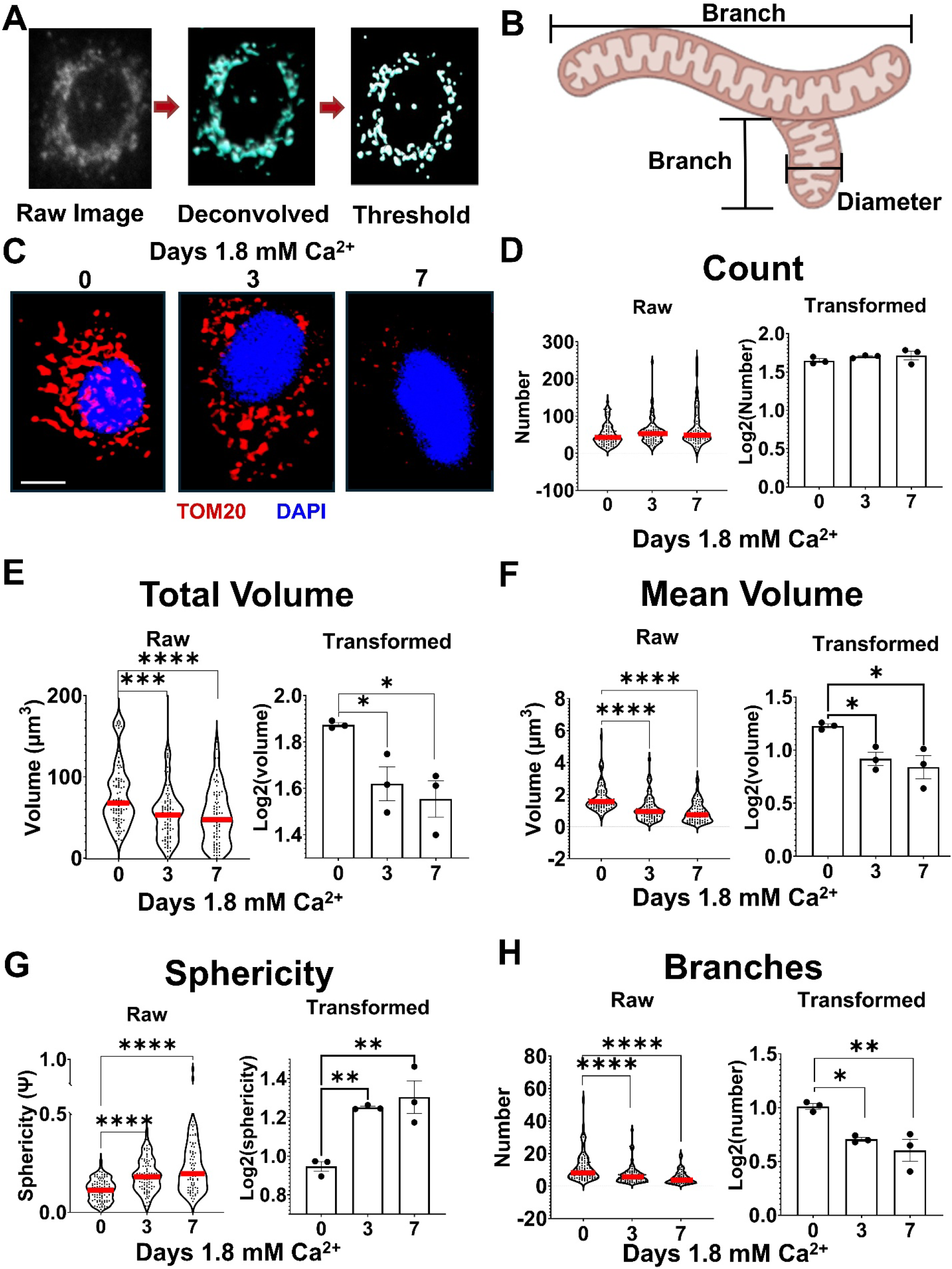
Esophageal epithelial differentiation is accompanied by mitochondrial network remodeling and fragmentation. (**A**) Representative image-processing workflow used for mitochondrial segmentation and morphometric analysis. (**B**) Schematic illustrating the mitochondrial parameters quantified. (**C**) Representative TOM20 immunofluorescence images of EPC2-hTERT cells following 0, 3, or 7 days of calcium-induced differentiation. (**D-H**) Automated three-dimensional quantification of mitochondrial number **(D**), total mitochondrial volume (**E**), mean mitochondrial volume (**F**), sphericity (**G**), and number of branches per mitochondrion (**H**), respectively. Raw single-cell measurements are shown on the left; each dot represents an individual cell; red line indicates mean. Data were obtained from three independent experiments (25 cells per trial, 75 cells total per treatment group). Trial-level transformed values are shown on the right; each dot represents an independent experiment; *p<0.05; **p<0.01; ***p<0.001; ****p<0.0001 by lognormal Brown-Forsythe and Welch’s ANOVA tests (raw data) or one-way ANOVA with Tukey’s post hoc test. Data are presented as log-transformed mean ± SEM of 3 independent trials.

### BNIP3L/NIX is the mitochondrial clearance-associated factor most consistently associated with esophageal epithelial differentiation

The reduction and fragmentation of mitochondria during epithelial differentiation suggested that regulated mitochondrial clearance may contribute to this process. Because mitochondrial fragmentation can facilitate mitochondrial elimination, we next evaluated the expression of several established mitophagy-associated factors during esophageal epithelial differentiation. We focused on *BNIP3L* (NIX), *BNIP3*, and *PARK2* (Parkin), which represent mitochondrial clearance pathways. Analysis of publicly available single-cell RNA-sequencing data from human esophageal epithelium showed that *BNIP3L* and *BNIP3* expression was preferentially associated with epithelial populations expressing the differentiation marker *IVL* (**Figure 5A, B**). In contrast, *PARK2* expression was low and restricted to a small population of esophageal epithelial cells (**Figure 5A, B**).

**Figure 5.**
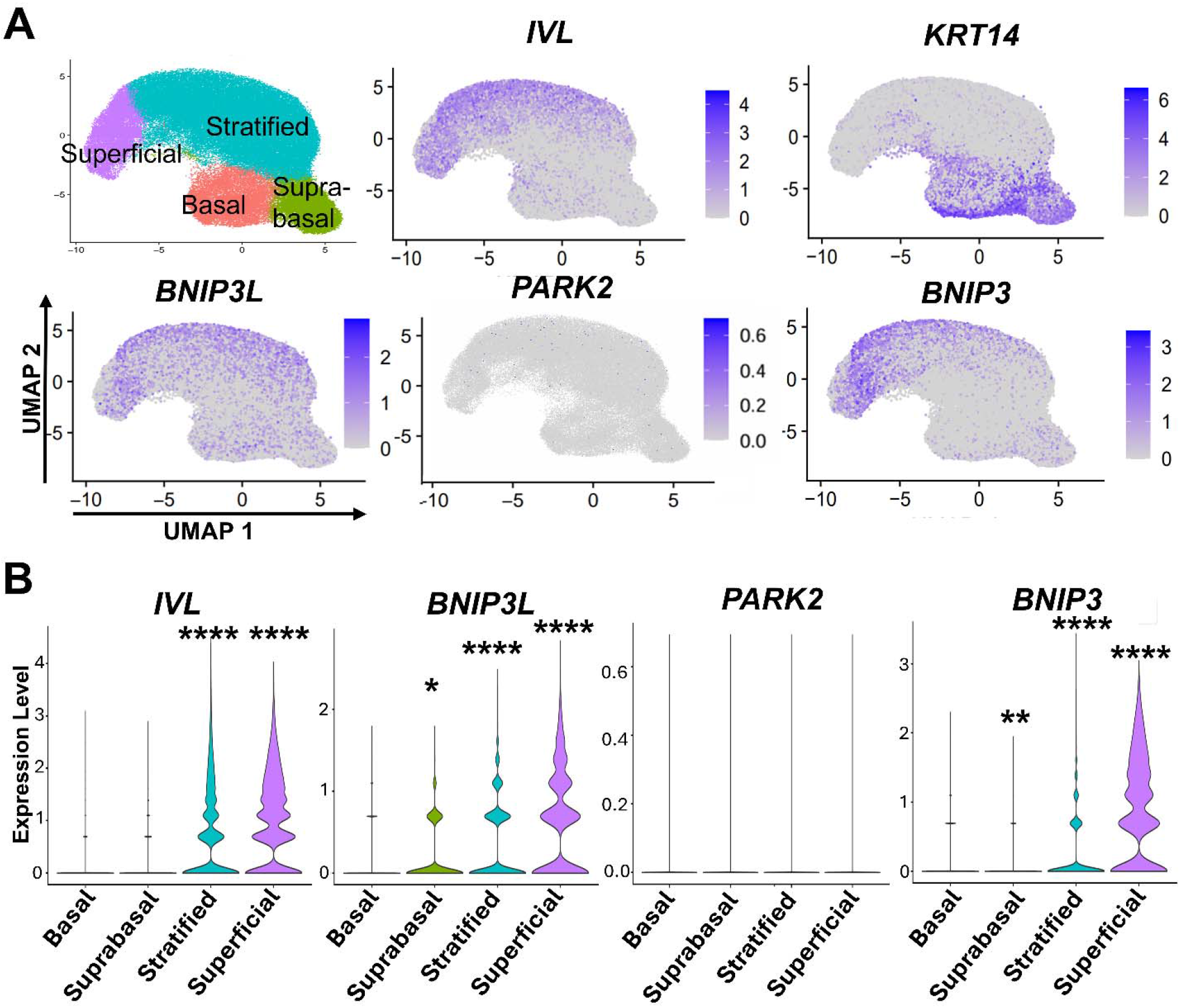
*BNIP3L* and *BNIP3* are enriched during differentiation in human esophageal epithelium. Expression of *IVL, KRT14, BNIP3L* (NIX)*, PARK2* (Parkin), and *BNIP3* in normal human esophageal epithelium from a published single-cell RNA-sequencing dataset (22). (**A**) Uniform Manifold Approximation and Projection (UMAP) plots with purple indicating enrichment. (**B**) Violin plots showing expression across epithelial cell compartments. Data include 6 independent human subjects, comprising n=89489 total cells combined from all subjects (Basal, n=10000; Suprabasal, n=10870; Stratified, n=59469; Superficial, n=9150). Statistical significance was determined by Wilcoxon rank-sum test comparing each cluster to the basal compartment; *p<0.05; **p<0.01; ****p<0.0001.

We also assessed EPC2-hTERT cells in an ALI culture, which generates a 3D stratified epithelium (24). In this model system, increased *IVL* and reduced *KRT14* expression supported EPC2-hTERT cell differentiation (**Figure 6A**). *BNIP3L* upregulation was also found in EPC2-hTERT ALI cultures, whereas *PARK2* and *BNIP3* expression was reduced (**Figure 6A**). We continued to examine mitophagy mediator expression during calcium- and confluence-induced differentiation of EPC2-hTERT cells *in vitro*. As expected, *IVL* increased in both model systems (**Figure 6B, C**). *BNIP3L* expression also increased significantly during differentiation in these model systems, while *PARK2* and *BNIP3* showed less pronounced changes (**Figure 6B, C**). Collectively, these data identified *BNIP3L* as being more consistently associated with esophageal epithelial differentiation than *PARK2* or *BNIP3* across multiple experimental systems.

**Figure 6.**
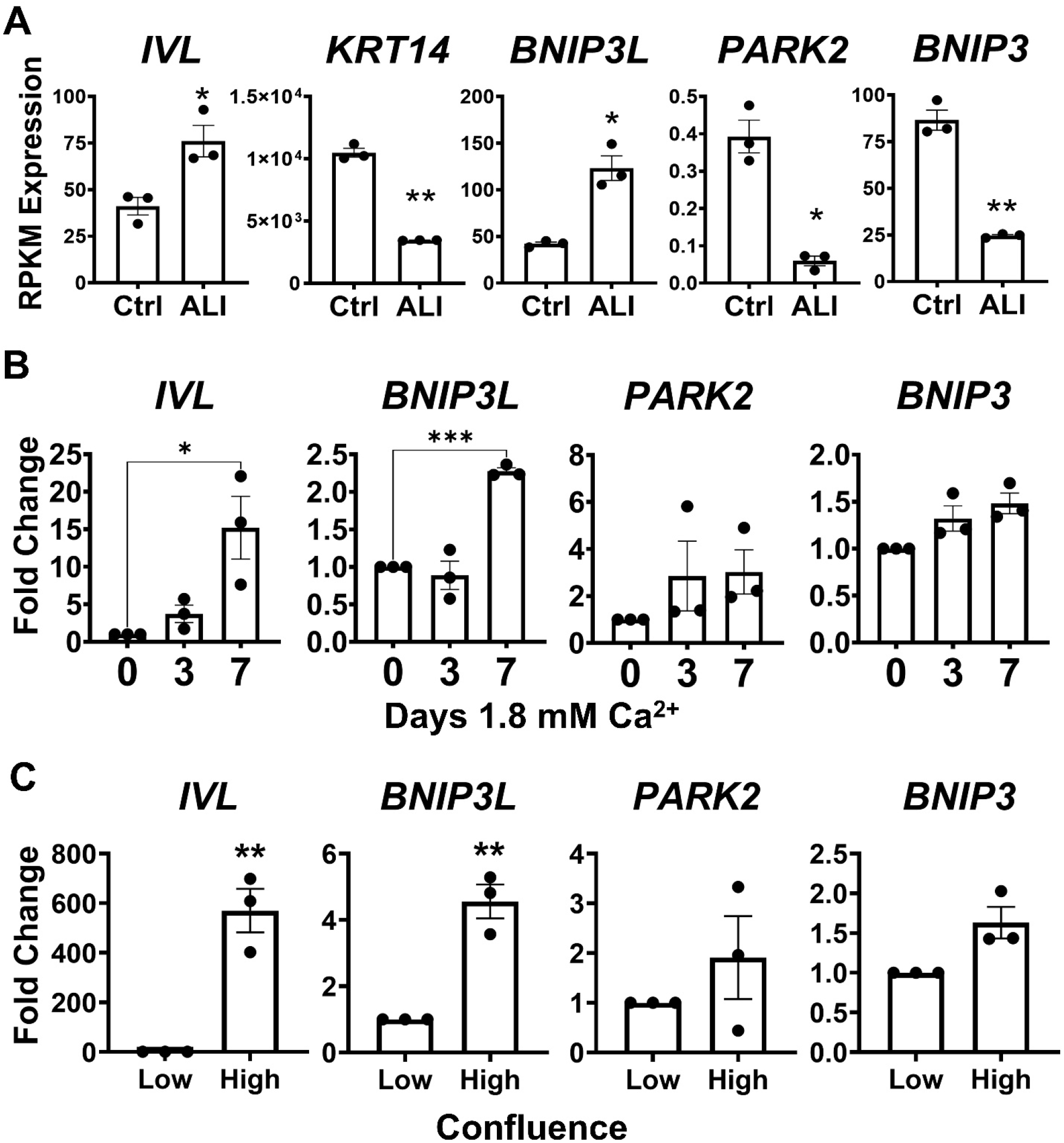
*BNIP3L* is induced in differentiation models of esophageal epithelial cells. (**A**) Quantification of *IVL, KRT14, BNIP3L* (NIX)*, PARK2,* and *BNIP3* RNA expression in control and air-liquid interface (ALI)-differentiated esophageal epithelial cultures. Data were obtained from EGID Express (25–27). Each dot represents an independent experiment. n=3; *p<0.05; **p<0.01 by Welch’s t-test. (**B**) EPC2-hTERT cells were treated with 1.8 mM Ca²⁺ for 0, 3, or 7 days to induce epithelial differentiation. Relative expression of *IVL*, *BNIP3L* (NIX), *PARK2* (Parkin), and *BNIP3* were measured by qRT-PCR. Each dot represents an independent experiment; n=3; *p<0.05; ***p<0.001 by one-way ANOVA with Tukey’s post hoc test. (**C**) EPC2-hTERT cells were cultured at low or high confluence. Relative expression of *IVL*, *BNIP3L* (NIX), *PARK2* (Parkin), and *BNIP3* was measured by qRT-PCR. Each dot represents an independent experiment; n=3; **p<0.01 by Welch’s t-test. Data are presented as mean± SEM of 3 independent trials.

### NIX is induced during esophageal epithelial differentiation *in vitro* and is associated with mitochondria

To determine whether NIX is recruited to the mitochondrial network during differentiation, we examined EPC2-hTERT cells via confocal microscopy. Upon calcium induction, NIX expression increased and was colocalized with TOM20-labeled mitochondria (**Figure 7A**). Quantitative analysis demonstrated a progressive reduction in TOM20 signal accompanied by increased NIX signal and an increased NIX-to-TOM20 ratio during differentiation (**Figure 7B-D**). Together, these findings demonstrate that NIX is induced at both the transcript and protein levels during esophageal epithelial differentiation and becomes increasingly associated with the mitochondrial compartment as mitochondrial abundance declines. Notably, immunocytochemistry and immunoblotting did not detect Parkin protein in EPC2-hTERT cells maintained under either undifferentiated or calcium-induced differentiation conditions. These findings further support NIX as the prominent mitochondrial clearance-associated factor induced during esophageal epithelial differentiation.

**Figure 7.**
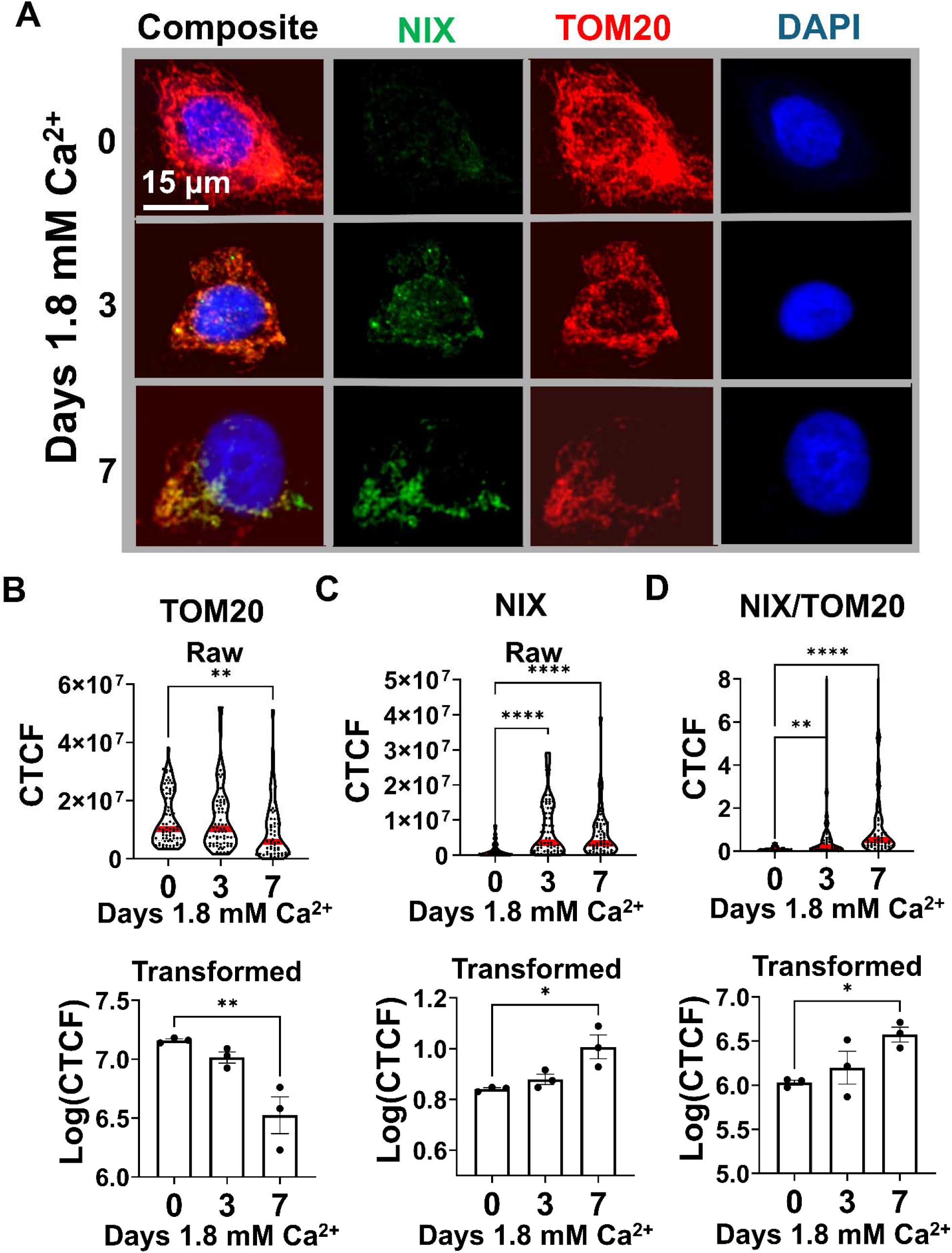
NIX is induced and associated with mitochondria during esophageal epithelial differentiation. EPC2-hTERT cells were treated with 1.8 mM Ca²⁺ for 0, 3, or 7 days to induce epithelial differentiation. (**A**) Representative confocal images of EPC2-hTERT cells stained for TOM20 (mitochondria; red), NIX (green), and DAPI (nuclei; blue) following treatment with 1.8 mM Ca²⁺ for 0, 3, or 7 days. Scale bar, 15 μm. (**B-D**) Quantification of fluorescence for TOM20 (**B**), NIX (**C**), and (**D**) NIX/TOM20 ratio. Raw single-cell measurements are shown above the transformed data; each dot represents an individual cell; the red line indicates the mean. Data were obtained from three independent experiments (25 cells per trial, 75 cells total per treatment group). Trial-level transformed values are shown below raw data; each dot represents an independent experiment; *p<0.05; **p<0.01; ****p<0.0001 by lognormal Brown-Forsythe and Welch’s ANOVA tests (raw data) or one-way ANOVA with Tukey’s post hoc test (transformed data). Log-transformed data are presented as mean ± SEM of 3 independent trials.

### NIX is required for differentiation-associated mitochondrial clearance and efficient esophageal epithelial differentiation

To determine whether NIX is functionally required for differentiation-associated mitochondrial depletion, we transfected EPC2-hTERT cells with either an NT control siRNA or an siRNA targeting *BNIP3L* and then maintained them in low-calcium or differentiation-inducing high-calcium medium (**Figure 8A**). We confirmed efficient *BNIP3L* depletion by qRT-PCR under both conditions. In control cells, high-calcium exposure robustly induced *KRT13*. This response was significantly attenuated following NIX depletion, indicating that NIX is required for an efficient differentiation response (**Figure 8A**). Consistent with these findings, immunoblotting demonstrated robust induction of NIX and CK13 in control cells following calcium-induced differentiation, whereas NIX depletion markedly reduced CK13 induction (**Figure 8B-C**).

**Figure 8.**
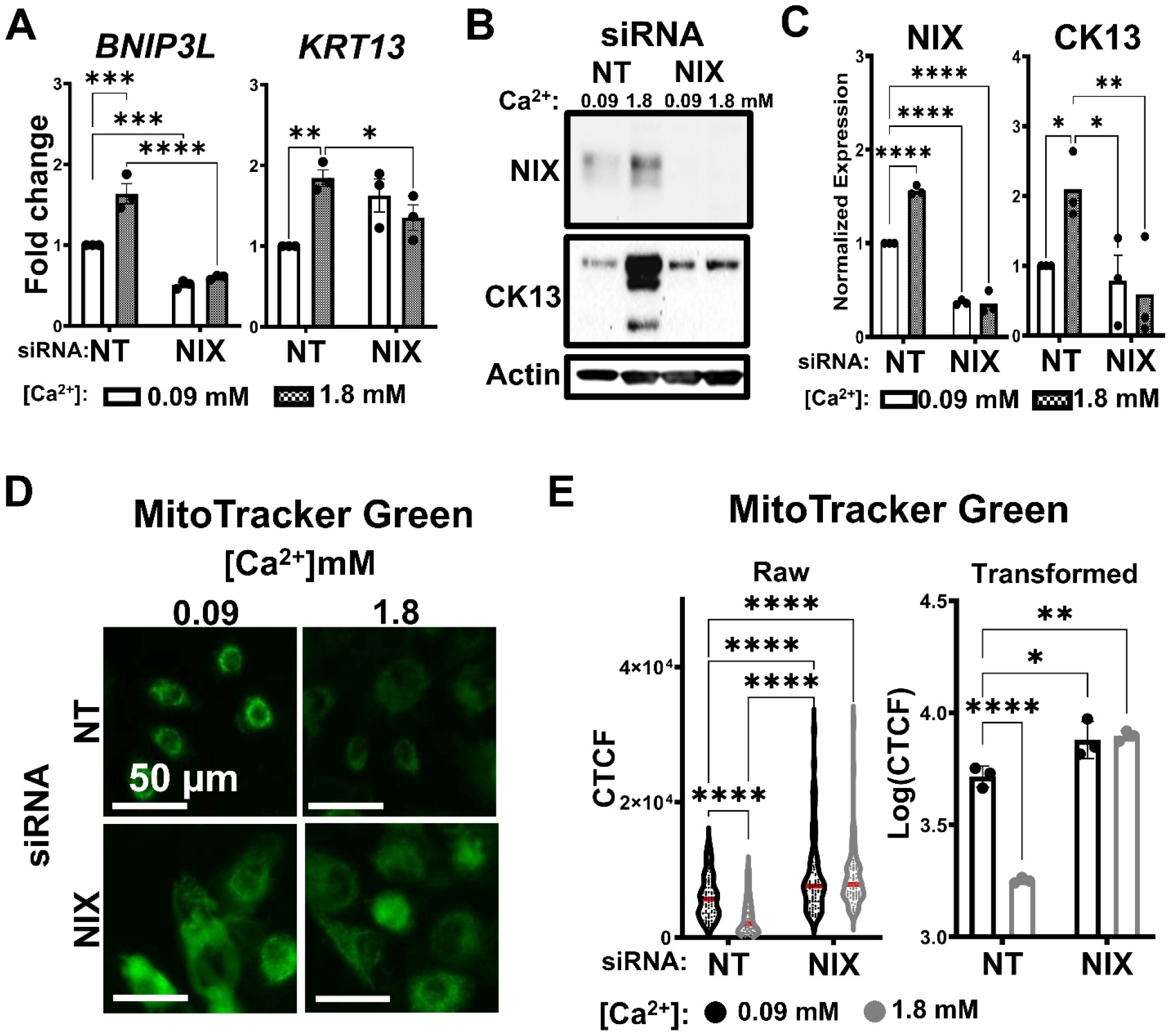
NIX is required for mitochondrial clearance and esophageal epithelial differentiation. EPC2-hTERT cells were transfected with a non-targeting control siRNA (siNT) or siRNA targeting NIX/*BNIP3L* (siNIX) and cultured in low-calcium (0.09 mM) or high-calcium (1.8 mM) medium. (**A**) Relative expression of *BNIP3L* and *KRT13* measured by qRT-PCR following NIX knockdown and calcium treatment. (**B**) Representative immunoblots of NIX and CK13 following NIX knockdown and calcium treatment. Actin was used as a loading control. (**C**) Densitometric quantification of NIX and CK13 protein expression from 3 independent trials. (**D**) Representative MitoTracker Green staining. (**E**) Quantification of MitoTracker Green fluorescence, shown as raw corrected total cell fluorescence (CTCF) and log-transformed values. Scale bars, 15 μm. Data were obtained from three independent experiments (25 cells per trial, 75 cells total per treatment group). Data are presented as mean ± SEM of three independent experiments. *p<0.05; **p<0.01; ***p<0.001; ****p<0.0001 by two-way ANOVA followed by Tukey’s post hoc test.

To determine whether NIX is required for differentiation-induced mitochondrial depletion, we evaluated mitochondrial mass following NIX knockdown. While high calcium exposure successfully triggered mitochondrial clearance in control cells, NIX-depleted cells failed to reduce their mitochondrial mass (**Figure 8D-E**). Furthermore, NIX depletion increased mitochondrial abundance even under low-calcium conditions (**Figure 8D-E**). These findings demonstrate that loss of NIX impairs both differentiation-associated mitochondrial depletion and epithelial differentiation, supporting a functional role for NIX in coordinating mitochondrial remodeling with acquisition of the differentiated phenotype.

### NIX expression coincides with mitochondrial clearance in the differentiating human esophageal epithelium

While our *in vitro* models demonstrate that NIX is required for mitochondrial depletion during differentiation, its spatial distribution in native tissue remains uncharacterized. Therefore, we sought to determine whether the observed phenotype *in vitro* is recapitulated in the human esophageal epithelium *in situ*. IHC analysis of normal human esophageal tissue demonstrated that NIX expression was detectable in the basal compartment, increased across the broader basal-to-suprabasal region, and subsequently declined toward the superficial epithelium (**Figure 9A**). In contrast, TOM20 expression was highest in basal cells and progressively decreased along the differentiation gradient (**Figure 9B**).

**Figure 9.**
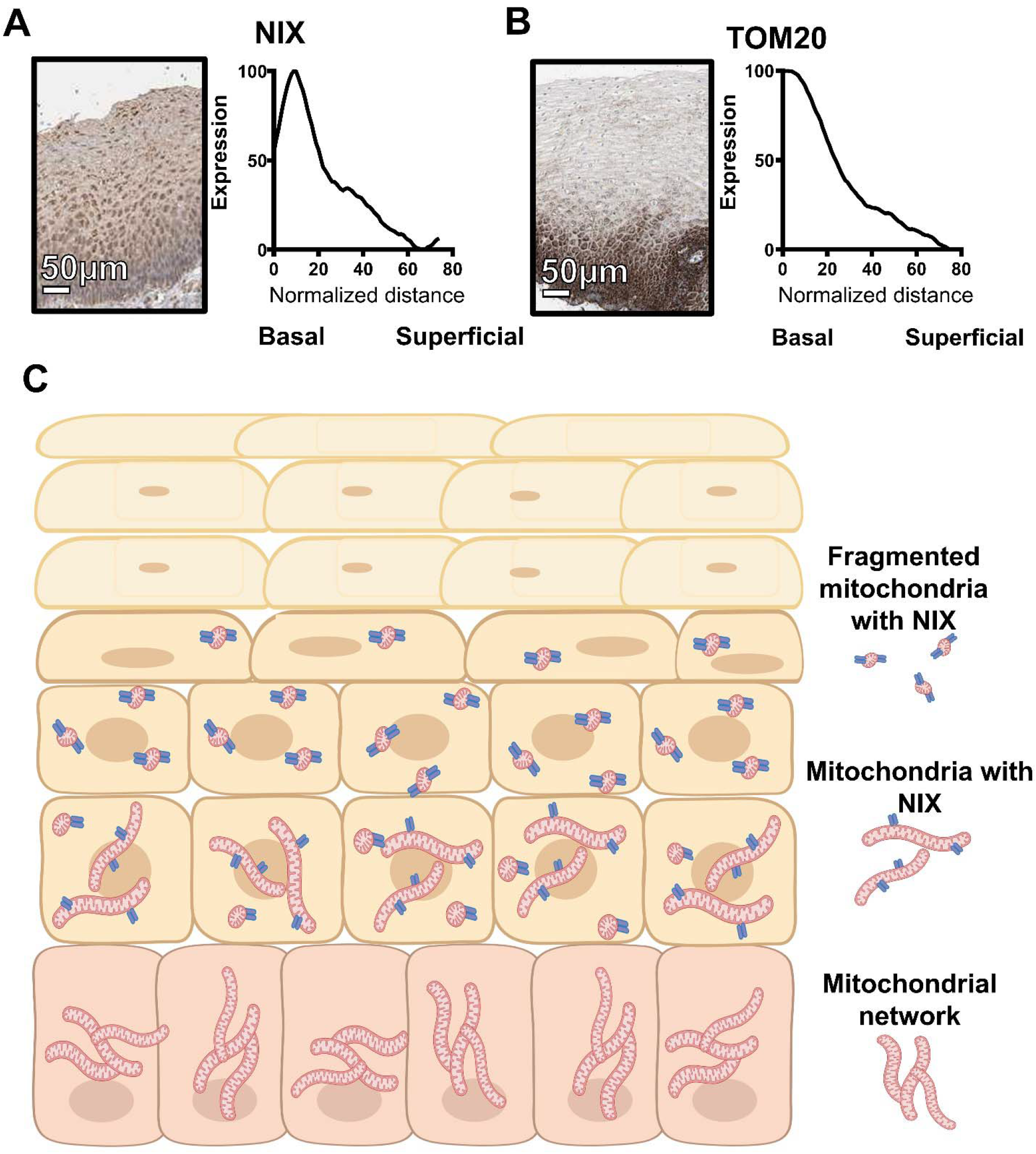
NIX is associated with the differentiation-associated mitochondrial gradient in human esophageal epithelium. (**A, B**) Immunohistochemical (IHC) staining for NIX (**A**) and TOM20 (**B**) in human esophageal squamous epithelium from the Human Protein Atlas was evaluated across the epithelium from basal to superficial. Scale bars, 50 μm. Line graphs represent staining patterns from 4 independent human subjects. (**C**) Proposed model of mitochondrial remodeling during esophageal epithelial differentiation. Basal cells exhibit high mitochondrial abundance and relatively low NIX expression. During progression through the basal-to-suprabasal compartment, NIX expression increases as the mitochondrial network undergoes fragmentation and mitochondrial abundance decreases. In the superficial epithelium, both NIX expression and mitochondrial abundance are low.

Together with our *in vitro* findings, these observations support a model in which NIX induction accompanies mitochondrial depletion during esophageal epithelial differentiation (**Figure 9C**). Specifically, we propose that basal cells exhibit high mitochondrial abundance with relatively low NIX expression. As cells transition through the basal-to-suprabasal compartment, the mitochondrial network undergoes remodeling, including fragmentation, accompanied by increased NIX expression and mitochondrial depletion. In the most superficial epithelium, both NIX expression and mitochondrial abundance are low.

### *BNIP3L*/NIX is suppressed in active EoE and restored following clinical remission

Lastly, to determine whether this differentiation-associated mitochondrial clearance program is disrupted in esophageal disease, we examined eosinophilic esophagitis (EoE), a condition characterized by impaired epithelial differentiation (10, 24). To validate the clinical relevance of our findings, we analyzed a published transcriptomic dataset of normal controls and patients with active EoE before and after fluticasone treatment (25–27). Compared with histologically normal controls, active EoE was characterized by reduced expression of the differentiation marker *IVL* and *BNIP3L*, coincident with increased expression of the mitochondrial marker *TIM23* (**Figure 10A**). To determine if these transcriptional defects are reversible upon resolution of inflammation, we evaluated the post-treatment cohort. Among patients who achieved remission after fluticasone treatment, *IVL* expression trended toward recovery, whereas *TIM23* expression trended toward reduction compared with active disease. Notably, *BNIP3L* expression was significantly increased following fluticasone-associated remission compared with active EoE (**Figure 10A**), suggesting that the suppression of this mitochondrial clearance program is associated with active disease and can be blunted by therapeutic intervention.

**Figure 10.**
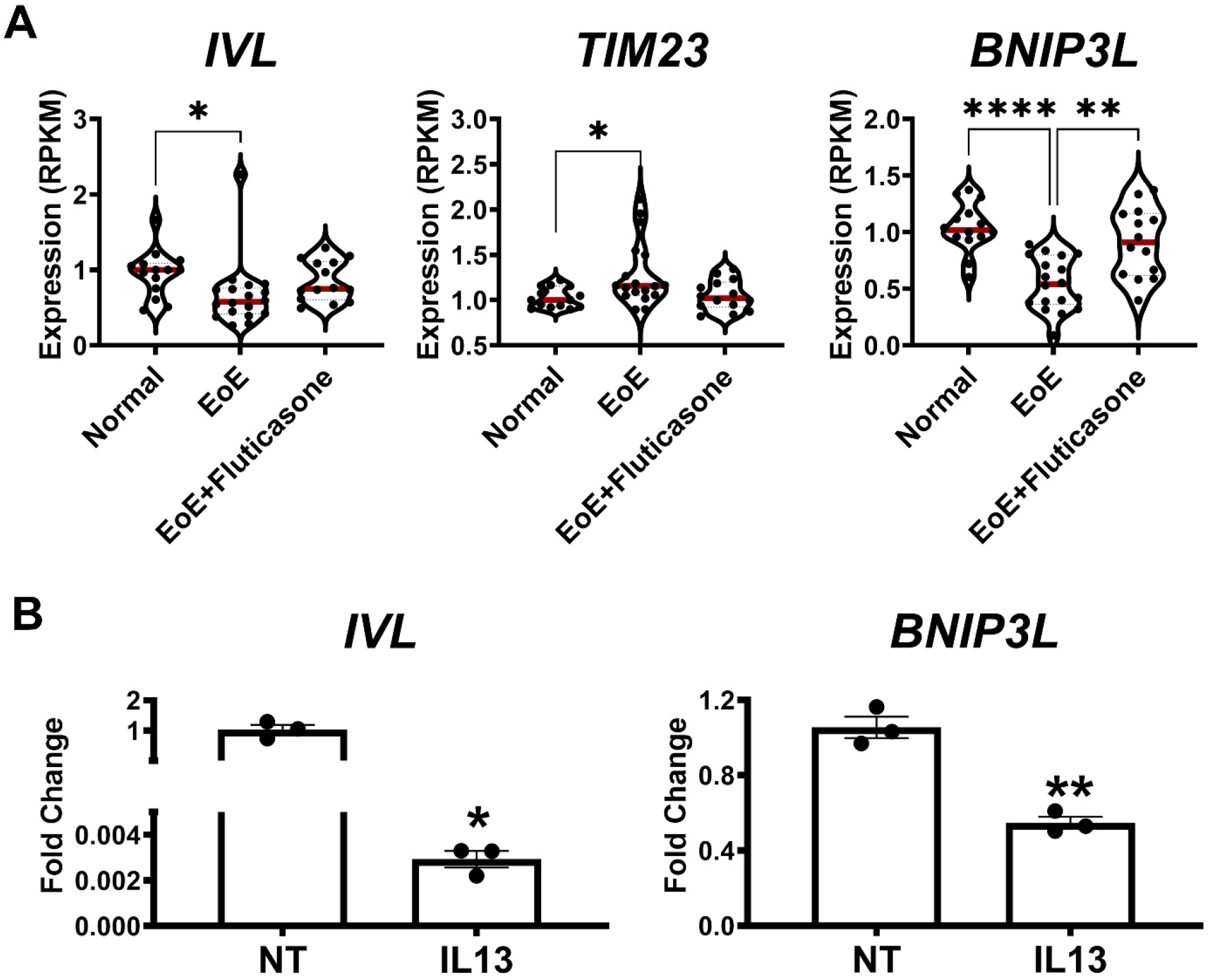
*BNIP3L*/NIX is suppressed in active EoE and partially restored following clinical remission. (**A**) Expression of the differentiation marker *IVL*, mitochondrial marker *TIM23*, and *BNIP3L* in esophageal epithelium from histologically normal controls (n=14), patients with active EoE (n=18), and patients with EoE who achieved remission following fluticasone treatment (n=14). Data were obtained from EGIDExpress. Each dot represents an independent human subject; red line represents mean; *p<0.05; **p<0.01; ****p<0.0001 by lognormal Brown-Forsythe and Welch’s ANOVA tests. (**B**) EPC2-hTERT cells were cultured in the presence or absence of IL-13 (10 ng/mL) for 7 days, then assessed for expression of *IVL* and *BNIP3L* by qRT-PCR. Each dot represents an independent experiment; n=3; *p<0.05; **p<0.01 by Welch’s t-test. Data are presented as mean ± SEM of 3 independent trials.

These findings suggested that inflammatory signals associated with EoE may suppress the NIX-associated mitochondrial clearance program. We therefore examined the effect of IL-13, a cytokine implicated in EoE pathogenesis and epithelial remodeling (25, 28), on EPC2-hTERT cells. Consistent with the findings in EoE tissue, IL-13 significantly reduced expression of both *IVL* and *BNIP3L* (**Figure 10B**). Thus, IL-13 exposure is sufficient to suppress both epithelial differentiation and expression of the NIX-associated mitochondrial clearance factor.

Together, these findings suggest that *BNIP3L*/NIX expression is reduced in active EoE and restored with clinical remission, linking inflammatory disease activity to disruption of a mitochondrial clearance-associated differentiation program. The suppression of *BNIP3L* in active disease, its restoration following clinical remission, and its reduction following IL-13 exposure suggest that inflammatory signaling may contribute to mitochondrial accumulation and impaired epithelial differentiation in EoE through suppression of NIX-associated mitochondrial clearance.

## Discussion

In this study, we identify mitochondrial remodeling as a previously unrecognized feature of homeostatic esophageal epithelial differentiation. Normal human esophageal epithelium exhibited a spatial gradient of mitochondrial abundance, with mitochondria enriched in the basal compartment and progressively depleted toward the superficial epithelium. This pattern was recapitulated in two independent *in vitro* models of differentiation, in which mitochondrial abundance decreased as differentiation markers increased. Our recent work established that autophagy is important for maintaining the stemness and self-renewal capacity of esophageal basal cells (29). Additionally, our previous studies linked Parkin-mediated mitophagy to epithelial-mesenchymal transition and cancer stem cell properties in esophageal cancer (30). Together, these observations suggest that autophagy and mitochondrial quality control are integrated with the regulation of esophageal epithelial cell state, with mitochondrial remodeling potentially contributing to the transition from self-renewal to differentiation.

Our findings further suggest that mitochondrial reduction contributes to, rather than merely accompanies, esophageal epithelial differentiation. Experimental depletion of TFAM reduced mitochondrial abundance and increased differentiation markers in the absence of an exogenous differentiation stimulus. Thus, reducing mitochondrial content was sufficient to promote features of the differentiated phenotype. This differs from studies in epidermal keratinocytes, where mitochondrial function and oxidative phosphorylation support the initiation of differentiation (7), emphasizing that the relationship between mitochondrial regulation and differentiation is context-dependent. Mitochondrial activity may support the energetic and biosynthetic requirements of proliferating basal cells, whereas the subsequent reduction in mitochondrial content may facilitate progression toward a more differentiated state.

We next sought to identify the mitochondrial clearance pathway associated with esophageal epithelial differentiation. Mitochondrial turnover can occur through multiple mechanisms, including PINK1/Parkin- and receptor-mediated pathways. *PARK2* expression was limited in human esophageal epithelial datasets, and Parkin protein was undetectable in EPC2-hTERT cells under basal or differentiation-inducing conditions. In contrast, *BNIP3L* was consistently associated with differentiated epithelial populations and was induced across multiple experimental differentiation models. These findings suggest that receptor-mediated mitochondrial quality-control pathways, particularly those involving NIX, may contribute to mitochondrial remodeling during esophageal epithelial differentiation. This does not exclude the role of Parkin in other epithelial states, which is consistent with our previous finding that Parkin-mediated mitophagy supports malignant progression in transformed esophageal epithelial cells (30).

Among the pathways examined, NIX emerged as the mitochondrial clearance-associated factor most consistently upregulated with esophageal epithelial differentiation. NIX increased during differentiation, became increasingly associated with TOM20-positive mitochondria as mitochondrial abundance declined, and was required for differentiation-associated mitochondrial depletion. NIX depletion also increased mitochondrial abundance under basal conditions, suggesting a role in mitochondrial turnover beyond differentiation. Importantly, our findings do not establish whether NIX promotes differentiation specifically through mitochondrial remodeling or through additional functions. NIX has also been implicated in apoptosis, endoplasmic reticulum function, and metabolic regulation (31, 32), which could independently influence the epithelial cell state. Determining whether the effects of NIX depletion on differentiation can be attributed specifically to its effects on mitochondrial abundance will therefore be an important direction for future studies.

The spatial distribution of NIX in human esophageal epithelium further supports a relationship between NIX expression and mitochondrial remodeling during differentiation. NIX expression increased from the basal compartment into the broader basal-to-suprabasal region before declining toward the superficial epithelium, whereas TOM20 progressively decreased along the differentiation gradient. Together with the temporal changes observed *in vitro*, this spatial pattern is consistent with transient NIX induction during the phase of differentiation associated with active mitochondrial remodeling. Although this interpretation remains to be directly tested, the relationship between NIX and mitochondrial abundance in human tissue is consistent with the model established by our *in vitro* studies.

Our analysis of human EoE samples further supported the relevance of this pathway to esophageal disease. Active EoE was associated with reduced *IVL* and *BNIP3L* expression and increased *TIM23* expression, whereas corticosteroid-associated remission increased *BNIP3L* and showed a trend toward reduced *TIM23*. These findings are consistent with an alteration of the differentiation-associated mitochondrial remodeling program in active EoE and suggest that this state is at least partially reversible with disease remission. We recently demonstrated increased mitochondrial mass in EoE (10), and the current findings provide a potential explanation for the persistence of mitochondrial content in epithelial cells that would normally undergo mitochondrial remodeling during differentiation. However, our findings do not establish that reduced NIX causes mitochondrial accumulation or impaired differentiation in EoE. Similarly, although IL-13 reduced both *BNIP3L* and *IVL* in EPC2-hTERT cells, these findings do not establish that IL-13 directly represses *BNIP3L* or that reduced NIX mediates its effects on mitochondrial abundance or differentiation. Restoration of *BNIP3L* following clinical response also raises the possibility that NIX-associated mitochondrial remodeling could serve as a biomarker of epithelial recovery, although this will require validation in longitudinal studies.

NIX regulation in EoE likely involves a broader network of inflammatory and metabolic signals. Hypoxia-inducible factor (HIF) signaling is of particular interest because HIF can regulate BNIP3 family members and has been implicated in epithelial-stress responses in the esophagus (11). At the same time, mitochondrial abnormalities in EoE are not uniform. Our previous work demonstrated increased mitochondrial mass (10), whereas other studies have reported reductions or functional impairment in specific aspects of mitochondrial activity (12). Thus, mitochondrial content, bioenergetic function, and turnover may be independently and dynamically regulated during EoE pathogenesis. More broadly, this concept may extend to epithelial aging, as our previous work identified mitochondrial dysfunction in aged esophageal epithelium. Whether age-associated changes in mitochondrial quality control alter NIX-dependent remodeling or influence the susceptibility to esophageal disease remains an important question.

In summary, our findings identify mitochondrial remodeling as an integral component of esophageal epithelial differentiation. Mitochondrial abundance progressively decreases as esophageal epithelial cells differentiate, accompanied by the remodeling and fragmentation of the mitochondrial network. Experimental mitochondrial depletion is sufficient to promote features of differentiation, whereas the loss of NIX prevents differentiation-associated mitochondrial depletion and impairs efficient differentiation. These findings identify NIX-associated mitochondrial remodeling as an important component of esophageal epithelial homeostasis and provide a potential mechanistic link between mitochondrial dysregulation and epithelial abnormalities in EoE. More broadly, our results highlight mitochondrial quality control as a previously underappreciated regulator of esophageal epithelial cell state.

## Abbreviations used in this paper

ALI: air-liquid interface
ANOVA: analysis of variance
BCL2: B cell lymphoma 2
BNIP3: BCL2/adenovirus E1B 19 kDa interacting protein 3
BNIP3L: BCL2/adenovirus E1B 19 kDa interacting protein 3 like
Ca²⁺: calcium
CK13: cytokeratin 13
CTCF: corrected total cell fluorescence
DOX: doxycycline
EGD: esophagogastroduodenoscopy
EoE: eosinophilic esophagitis
H&E: hematoxylin and eosin
HIF: hypoxia-inducible factor
IF: immunofluorescence
IHC: immunohistochemistry
IL: interleukin
IVL: involucrin
KRT13: cytokeratin 13
KRT14: cytokeratin 14
KSFM: keratinocyte serum-free medium
MTCO1: mitochondrially encoded cytochrome C oxidase 1
MTND3: mitochondrial NADH dehydrogenase subunit 3
NIX: Nip3-like protein X
NT: non-targeting
OXPHOS: oxidative phosphorylation
PARK2: Parkin
PBS: phosphate-buffered saline
qRT-PCR: quantitative reverse transcription polymerase chain reaction
ROI: region of interest
scRNA-seq: single-cell RNA sequencing
SEM: standard error of the mean
shRNA: short hairpin RNA
siRNA: small interfering RNA
TFAM: transcription factor A mitochondrial
TIM23: translocase of the inner mitochondrial membrane
TOM20: translocase of outer mitochondrial membrane 20
TUH: Temple University Hospital
UMAP: uniform manifold approximation and projection.

## Data Availability Statement

Data, analytic materials, and study materials will be made available upon reasonable request to Dr. Whelan.

## Author Contribution

N.S., M.R., A.B.M., Z.W.R., and K.A.W. conceived and designed the research. N.S., A.B., A.T., A.D.F., J.L.J., A.J.S., J.M.C., W.N-L., and K.Q.C. performed experiments. N.S., A.B., A.D.F., W.N-L., N.M.M., Z.W.R., and K.A.W. analyzed data. N.S. and K.A.W. interpreted data, prepared figures, and drafted the manuscript. N.S., A.B., A.T., A.D.F., J.L.J., A.J.S., J.M.C., W.N-L., N.M.M., A.J.K-S., K.Q.C., M.R., A.B.M., Z.W.R., and K.A.W. edited and revised the manuscript. N.S., A.B., A.D.F., J.L.J., A.J.S., J.M.C., W.N-L., A.T., N.M.M., A.J.K-S., K.Q.C., M.R., A.B.M., Z.W.R., and K.A.W. approved the final version.

