## Supplementary Figure 1 for "NIX-associated mitochondrial remodeling contributes to esophageal epithelial differentiation"

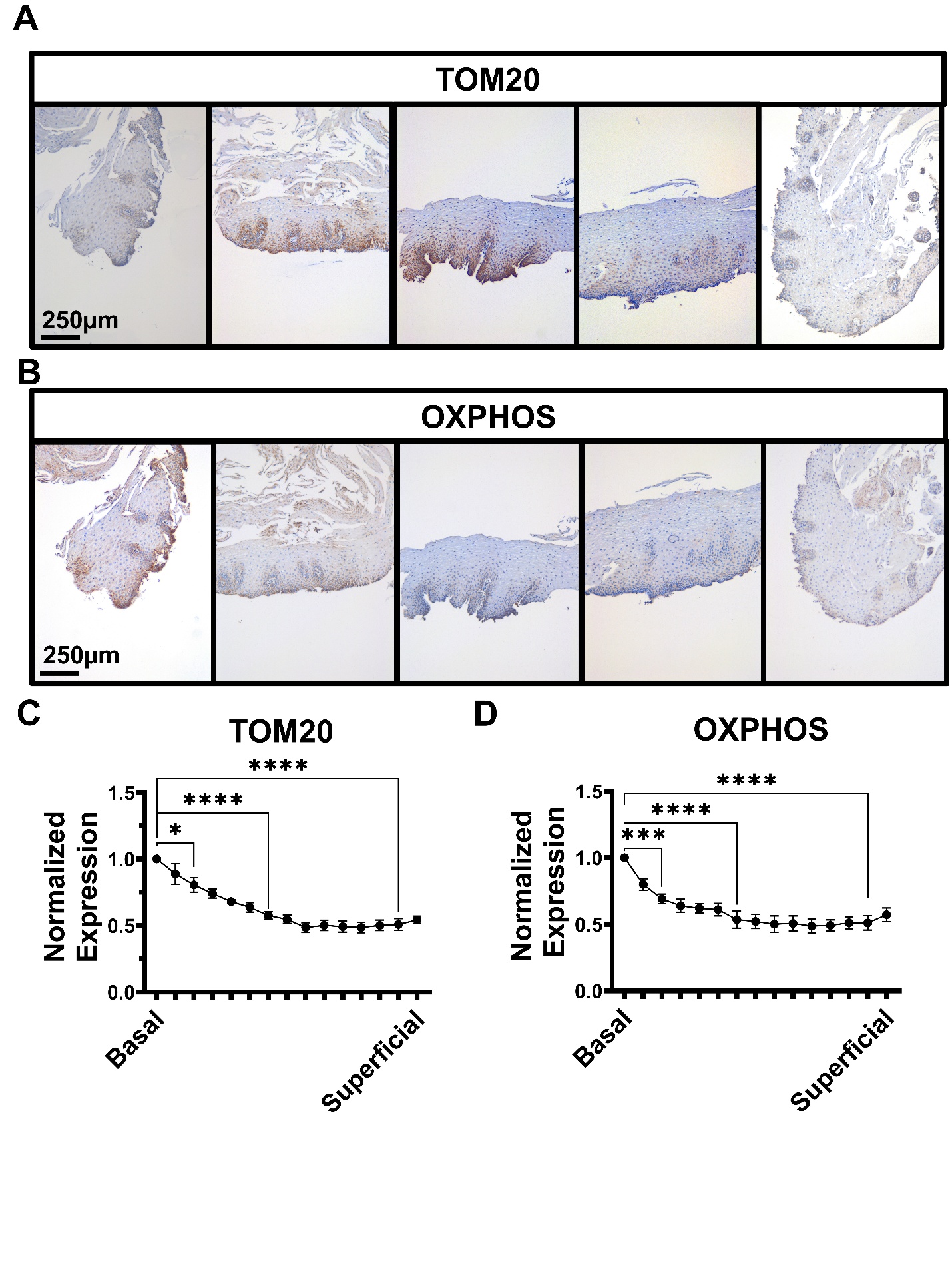

**Supplementary Figure 1:** **Mitochondrial marker expression is decreased across the basal-to-superficial axis in human esophageal epithelium.** (**A, B**) Representative immunohistochemistry staining images demonstrating the spatial distribution and expression patterns of (**A**) TOM20 and (**B**) OXPHOS across multiple representative tissue sections. Scale bars, 250 µm. (**C, D**) Quantification of normalized expression levels for (**C**) TOM20 and (**D**) OXPHOS across the epithelial stratification axis from the basal layer to the superficial layer. Data are presented as mean ± SEM from 5 human subjects, analyzed by one-way ANOVA followed by Tukey’s post hoc test; *p<0.05; ***p<0.001; ****p<0.0001.
